# Smooth Muscle Cells and Fibroblasts in the Proximal Thoracic Aorta Exhibit Minor Differences Between Embryonic Origins in Angiotensin II-driven Transcriptional Alterations

**DOI:** 10.1101/2025.01.23.610985

**Authors:** Sohei Ito, David B. Graf, Yuriko Katsumata, Jessica J. Moorleghen, Chen Zhang, Yanming Li, Scott A. LeMaire, Ying H. Shen, Hong S. Lu, Alan Daugherty, Hisashi Sawada

## Abstract

**Background:** Thoracic aortopathy is influenced by angiotensin II (AngII) and exhibits regional heterogeneity with the proximal region of the thoracic aorta being susceptible. Smooth muscle cells (SMCs) and selected fibroblasts in this region are derived from two embryonic origins: second heart field (SHF) and cardiac neural crest (CNC). While our previous study revealed a critical role of SHF-derived cells in AngII-mediated aortopathy formation, the contribution of CNC-derived cells remains unclear.

**Methods:** Mef2c-Cre R26R^mT/mG^ mice were infused with AngII (1,000 ng/kg/min). Proximal thoracic aortas were harvested at baseline or after 3 days of infusion, representing the prepathological phase. Cells were sorted by origins using mGFP (SHF-derived) and mTomato (other origins, nSHF-derived) signals, respectively. After sorting cells by origin, single-cell RNA sequencing was performed and analyzed.

**Results:** Short-term AngII infusion induced significant transcriptomic changes in both SHF- and nSHF-derived SMCs, but differences between origins were modest. Fibroblast transcriptomes also underwent notable changes by AngII infusion, but differences between SHF and nSHF origins remained modest. Interestingly, AngII infusion resulted in the emergence of a new fibroblast sub-population. Several molecules related to the extracellular matrix, such as *Eln* and *Col3a1*, were downregulated in SHF-derived fibroblasts compared to nSHF-derived fibroblasts in the new subcluster.

**Conclusion:** Fibroblasts in the new subcluster exhibited lineage-specific differences in extracellular matrix-related genes; however, overall transcriptomic differences between origins in SMCs and fibroblasts in response to AngII were modest in the pre-pathological phase of AngII-induced thoracic aortopathy.

**GRAPHIC ABSTRACT:** 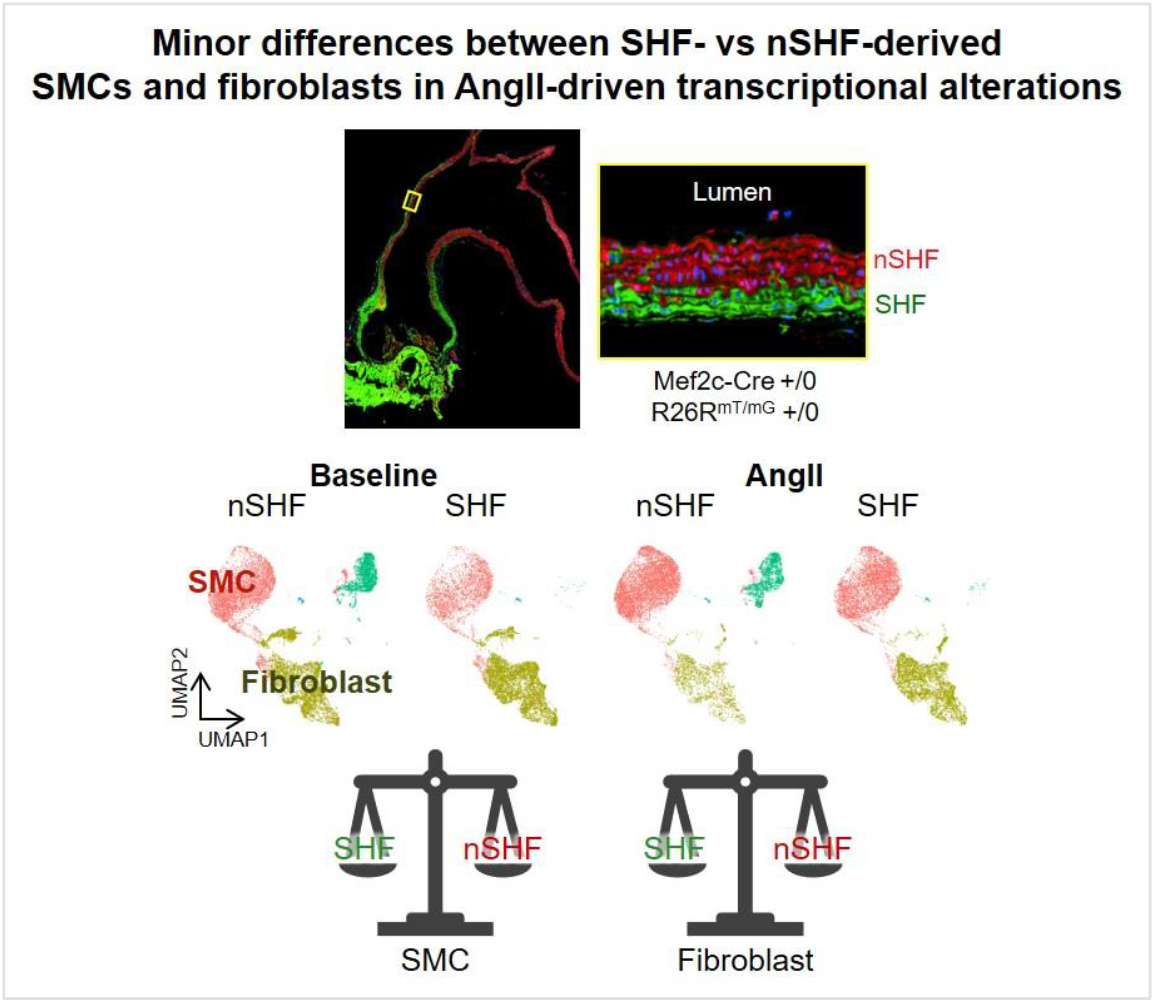

## INTRODUCTION

Thoracic aortopathy is a spectrum of life-threatening diseases characterized by aortic aneurysm, dissection, or rupture.^1,2^ Despite its significant burden on public health, thoracic aortopathy lacks effective pharmacological therapy to inhibit disease initiation and development. Thus, understanding the mechanisms underlying thoracic aortopathy formation is crucial for developing effective treatments. In thoracic aortopathy, aneurysms develop frequently in the aortic root and ascending aorta.^3,4^ Aortic dissection occurs preferentially in the outer third of the media.^5^ Vascular wall thickening, a histological hallmark of thoracic aortopathy, is mainly observed in the outer media of the aortic wall.^6-11^ These features suggest that the outer media of the proximal thoracic aorta, including the aortic root and ascending aorta, is especially susceptible to disease development. However, mechanisms underlying these regional heterogeneities have not been defined.

Smooth muscle cells (SMCs), the predominant cell type in the aortic media, exert a pivotal role in the pathophysiology of vascular diseases, including thoracic aortopathy.^12^ In the thoracic aortopathy-prone regions, aortic root and ascending aorta, SMCs are derived from two distinct embryonic origins: cardiac neural crest (CNC) and second heart field (SHF).^13,14^ SMCs from these two origins reside in spatially distinct domains. CNC-derived SMCs extend from the aortic root to the distal end of the aortic arch, whereas SHF-derived SMCs are distributed from the aortic root to just proximal to the innominate artery. Moreover, in the ascending aorta, CNC-derived SMCs predominantly populate the inner medial layers in the outer curvature and form a mosaic pattern in the inner curvature, whereas SHF-derived SMCs are localized to the outer medial layers. Thus, SHF-derived SMCs are predominant in the regions susceptible to thoracic aortopathy, and functional differences between these two SMC lineages could determine mechanisms underlying thoracic aortopathy.

Angiotensin II (AngII), a potent vasoactive peptide, drives a spectrum of pathological processes in thoracic aortopathy. In mice, AngII infusion induces thoracic aortopathy in the ascending aorta,^15^ closely replicating the regional heterogeneity observed commonly in human thoracic aortopathy. Our previous study identified SHF-derived cells as a key contributor to AngII-mediated thoracic aortopathy.^16^ Deletion of low-density lipoprotein receptor-related protein 1 (LRP1) in SHF-derived cells exacerbated AngII-induced thoracic aortopathy. Single-cell RNA sequencing (scRNAseq) combined with lineage tracing further revealed that AngII infusion induces significant transcriptomic alterations in SHF-derived cells, particularly in genes associated with the extracellular matrix and the transforming growth factor β (TGF-β) signaling pathway, prior to thoracic aortopathy formation. Deletion of TGF-β type 2 receptor (TGFBR2) in SHF-derived cells results in embryonic lethality accompanied by outflow tract dilatation and retroperitoneal hemorrhage, suggesting the importance of TGF-β signaling in SHF-derived cells in aortic integrity. Despite these significant findings regarding SHF-derived cells, the role of CNC-derived cells in AngII-mediated thoracic aortoatphy formation is largely unknown.

In this study, we determined aortic phenotypes in mice with LRP1 or TGFBR2 in CNC-derived cells. In addition, using scRNAseq analyses with SHF-lineage tracing, we also determined whether CNC-derived cells exhibit distinct transcriptomic alterations in response to AngII from SHF-derived cells prior to thoracic aortopathy formation.

## METHODS

### Mice

ROSA26R^LacZ^ (#003474), ROSA26R^mTmG^ (#007676), *Wnt1*-*Cre* (#022137), *Lrp1* floxed (#012604), and *Tgfbr2* floxed (#012603) mice were purchased from The Jackson Laboratory. Mef2c-*Cre* mice (#030262) were purchased from the Mutant Mouse Resource and Research Center. *Wnt1*- or *Mef2c*-*Cre* male mice were bred to ROSA26R^LacZ^ female mice to trace CNC- or SHF-derived cells, respectively. *Mef2c*-*Cre* +/0 male mice were also bred to ROSA26R^mTmG^ female mice for scRNAseq analyses. To delete LRP1 in CNC-derived cells, *Lrp1* floxed female mice were bred with *Wnt1-Cre* mice. *Wnt1-Cre* male mice were also used for CNC-specific deletion of TGFBR2. Mice were fed a normal laboratory rodent diet and provided with drinking water. Mice were randomly assigned to study groups. All procedures were approved by the University of Kentucky’s IACUC.

### Embryonic Study

Embryonic studies were performed as described previously. Briefly, to investigate aortic malformation during the prenatal stage, fetuses were harvested from pregnant females on either E11.5 or E12.5. The morning following the detection of a vaginal plug in mated females was defined as E0.5 of gestation. Female breeders were euthanized via intraperitoneal injection of a ketamine/xylazine cocktail (90 and 10 mg/kg, respectively), followed by saline perfusion (8 mL) through the left ventricle. The abdominal cavity was opened, and fetuses were dissected carefully. Gross fetal morphology was documented using a dissection microscope with a high-resolution camera (SMZ800, DS-Ri1, Nikon). Cranial tissue was collected for genotyping. Embryos were then fixed in buffered formalin (10% wt/vol). After 24 hours, the chest wall, pericardium, and atriums were removed gently, and the gross morphology of the outflow tract was captured using a dissection microscope coupled with a high-resolution camera.

### Pump implantation

AngII (1,000 ng/kg/min, H-1705, Bachem) was infused via a subcutaneously implanted osmotic pump (Alzet model 2001) into male mice at 10 to 12 weeks of age, as described previously.^17^ Surgical staples were used to close incision sites. Post-operative pain was alleviated by application of topical LMX4 lidocaine cream (4% wt/wt, #0496-0882-15, Eloquest Healthcare Inc).

### X gal staining

Whole tissues were incubated with X-gal (V3941, Promega) and eosin B (E8764, Sigma). Briefly, tissues were fixed in paraformaldehyde (4% wt/vol), and incubated in buffer containing sodium phosphate (100 mM, pH 7.3), MgCl_2_ (2 mM), sodium deoxycholate (0.01% wt/vol) and NP40 (0.02% wt/vol) for 90 minutes. Subsequently, X-gal (1 mg/mL), potassium ferricyanide (5 mM), and potassium ferrocyanide (5 mM) were added to the buffer and samples were incubated overnight at room temperature. Whole tissues were post-fixed with 10% buffered formalin. Distribution data were validated by a researcher blinded to the study group information.

### Confocal microscopy

Aortic samples were fixed with paraformaldehyde (4% wt/vol) for 24 hours and sucrose (30% wt/vol) for 24 hours. Subsequently, aortic tissues were embedded into optimum cutting temperature (OCT) compound (#4583, Sakura Finetek) and cut into 10 μm sections. Aortic sections were washed with PBS for three times and mounted using an anti-fade reagent with DAPI (#ab104139, abcam). Fluorescent images were captured at 20x magnification using the tile image function by a Nikon AXR inverted confocal microscope.

### scRNAseq

Ascending aortic samples were harvested from *Mef2c-Cre* ROSA26R^mT/mG^ +/0 male mice at baseline (n=5) and after 3 days of AngII infusion (n=4). Periaortic tissues were removed and aortic tissues were cut into small pieces. Single-cell suspension was performed as described previously. Briefly, aortic samples were digested with an enzyme cocktail containing collagenase type II (3 mg/mL, #LS004176, Worthington), collagenase type XI (0.15 mg/mL, #C7657, Sigma-Aldrich), Hyaluronidase type I (0.24mg/mL, #H3506, Sigma-Aldrich), elastase (0.1875 mg/mL, #LS002290, Worthington) in Ca/Mg contained-HBSS (#14025092, Thermo Fisher Scientific) for 60 minutes at 37°C. Cells were then filtered through a 40 μm cell strainer (CLS431750-50EA, Sigma-Aldrich), centrifuged at 300 g for 10 minutes, and resuspended using cold HBSS (#14175095) with fetal bovine serum (5% vol/vol). Subsequently, cells were sorted based on mTomato and mGFP signals by flow cytometry (FACS Aria III, BD Biosciences).

mTomato and mGFP positive cells were dispensed separately onto the Chromium Controller (10x Genomics) and indexed single cell libraries were constructed by a Chromium Single Cell 3’ v3 Reagent Kit (10x Genomics). cDNA libraries were sequenced in a pair-end fashion on an Illumina NovaSeq 6000. Raw FASTQ data were aligned to the mouse transcriptome (mm10) by Cell Ranger 3.0 (baseline) or 5.0.1 (AngII).

### Statistical analysis for scRNAseq data

Seurat package (v4.3.0) and R (v4.1.0) were used to analyze scRNAseq data on R studio (2022.12.0).^18,19^ Four mapped unique molecular identifier (UMI) counts datasets from mTomato and mGFP positive cells (baseline-SHF, baseline-nSHF, AngII-SHF, and AngII-nSHF) were imported into R separately. Seurat objects for each of the UMI count datasets were built using the “CreateSeuratObject” function by the following criteria: ≥3 cells and ≥200 detected genes. Cells expressing less than 200 or more than 5,000 genes were filtered out for exclusion of non-cell or cell aggregates, respectively. Cells with more than 10% mitochondrial genes were also excluded. UMI counts were then normalized as follows: counts for each cell were divided by total counts, multiplied by 10,000, and transformed to a natural log. “FindIntegrationAnchors” and “IntegrateData” functions were used to remove batch effects and integrate the four normalized datasets. Uniform manifold approximation and projection (UMAP) dimensionality reduction to 20 dimensions for the first 30 principal components (PCs) was applied to identify cell clusters using the normalized- and scaled-UMI count data. “FindAllMarkers” and “FindConcervedMarkers” functions were used to identify conserved marker genes to determine cell types of each of the clusters. Differentially expressed genes were examined using a zero-inflated negative binomial regression model with two factors (embryonic origin and infusion) and their interaction from “zinbwave (1.16.0)” and “edgeR (v3.36.0)” R packages.^20,21^ Genes start with “Gm”, “Rp”, or “mt-” and end with “Rik” genes were excluded. FDR was calculated by “p.adjust” using “fdr” method. FDR < 0.05 was used as a threshold to identify differentially expressed genes. Gene ontology enrichment analyses for biological process were performed using “clusterProfiler (4.0.2)” R package.^22^ R codes and scRNAseq data were validated by a biostatistician.

### Data Availability

Raw and processed scRNAseq data are available at the Gene Expression Omnibus (GSE275132). Analyzed data have been attached as Supplemental Excel File.

## RESULTS

### CNC-specific LRP1 deletion induced embryonic lethality, and CNC-specific TGFBR2 deletion resulting in aortic deformities in neonates

We first determined the impact of LRP1 deletion in CNC-derived cells on AngII-induced thoracic aortopathy. To develop CNC-specific LRP1 deficient mice, *Lrp1* floxed mice were bred with *Wnt1-Cre* mice targeting cells derived from the CNC origin. As we reported previously,^16^ mice with LRP1 deletion in SHF-derived cells were vital and had normal development, but showed more severe AngII-induced thoracic aortopathy in adult mice, compared to their wild-type littermates. In contrast, LRP1 deletion in CNC-derived cells resulted in embryonic lethality. While fetuses appeared intact at E11.5, fetal resorption was observed at E12.5 (**Supplemental Figure 1A, B**). In situ imaging showed no discernable structural abnormalities were detected in the heart or outflow tract at E11.5 (**Supplemental Figure 1B**). X-gal staining of major organs revealed that *Wnt1-Cre* induced recombination in a broader range of organs compared to *Mef2c-Cre* (**Supplemental Figure 2A, B**), suggesting that the widespread distribution of *Wnt1*-driven cells contributes to embryonic lethality by LRP1-deletion using Wnt1-Cre. Given embryonic lethality, we concluded that LRP1 deletion using *Wnt1-Cre* is not optimal for investigating the role of CNC-derived cells in AngII-mediated thoracic aortopathy.

As an alternative approach, we next generated CNC-specific TGFBR2 deficient mice by crossing *Tgfbr2* floxed mice with *Wnt1*-*Cre* mice. Unlike LRP1 deletion, mice lacking TGFBR2 in CNC-derived cells were born alive. However, these neonates displayed calvarial defects (**Supplemental Figure 3A**). In addition, in situ imaging confirmed persistent truncus arteriosus in these mice (**Supplemental Figure 3B**). These results are consistent with previous studies reporting that TGFBR2 deletion in CNC-derived cells leads to calvarial defects, micrognathia, and persistent truncus arteriosus.^23-25^ Our previous study demonstrated that TGFBR2 deletion in SHF-derived cells results in embryonic lethality accompanied by outflow tract dilation and peritoneal hemorrhage. Together, these findings indicate both SHF- and CNC-derived cells are necessary for normal aortic development through TGF-β signaling. However, similar to LRP1 deletion in CNC-derived cells, we concluded that CNC-specific TGFBR2 deletion is not a viable model for determining the role of CNC-derived cells in AngII-mediated thoracic aortopathy.

### scRNAseq analyses using SHF-lineage tracing mice with short-term AngII infusion

Next, to determine whether CNC-derived cells display divergent transcriptomic alterations from SHF-derived SMCs in AngII-mediated thoracic aortopathy formation, scRNAseq was performed by pooling cells from ascending aortas of male *Mef2c-Cre* +/0 R26R^mTmG^ +/0 mice (**Figure 1A, B**).^16^ Aortic samples were harvested before and 3 days after AngII infusion. Samples exhibiting discernable aortopathy were excluded to focus on the initiation process of AngII-mediated thoracic aortopathy. In *Mef2c-Cre* +/0 R26R^mTmG^ +/0 mice, SHF-derived cells are traced by mGFP, whereas cells not derived from the SHF origin (nSHF) express mTomato (**Figure 1A**). Since SMCs in the ascending aorta are from either SHF or CNC origin, most non-SHF-derived SMCs are considered to be derived from the CNC. SHF- and nSHF-derived cells were sorted based on their respective fluorescent signals by FACS and sequenced separately. Data from SHF-derived cells were reported in our previous study investigating the role of SHF-derived cells. Since the present study aimed to identify transcriptomic differences between origins in response to AngII, data from nSHF-derived cells were integrated with published data from SHF-derived cells and analyzed by two-way ANOVA for the interaction between AngII infusion and SMC origins (**Figure 1B**).

**Figure 1.**
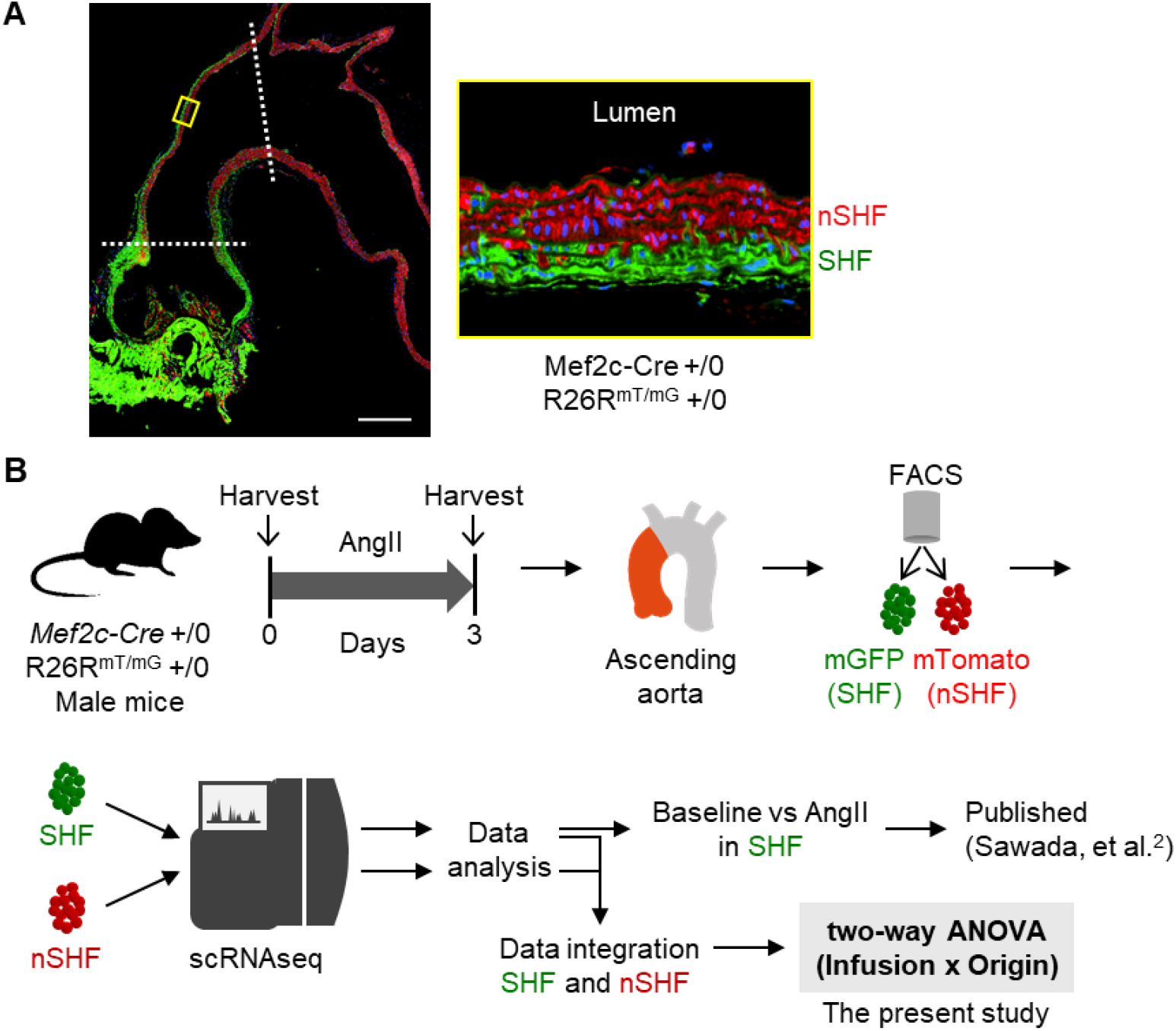
Workflow for scRNAseq analyses using Mef2c-Cre R26R^mTmG^ mice with short-term AngII infusion. **(A)** Representative confocal images of the thoracic aorta from male Mef2c-Cre +/0 R26R^mTmG^ +/0 mice. Scale bar = 500 μm. **(B)** Study workflow. Proximal thoracic aortas were harvested from male Mef2c-Cre +/0 R26R^mTmG^ +/0 mice at baseline and after 3 days of AngII infusion (n = 4–5/group). SHF-derived (SHF) and non-SHF-derived (nSHF) cells were separated via FACS based on mGFP and mTomato signals and sequenced independently. Read count data of SHF and nSHF cells at baseline and after 3 days of AngII infusion were integrated and analyzed by two-way ANOVA.

### Modest transcriptomic differences between SHF- and nSHF-derived SMCs in the prepathological phase of AngII-induced thoracic aortopathy

After removing batch effects, the four normalized datasets (nSHF-baseline, SHF-baseline, nSHF-AngII, SHF-AngII) were integrated properly (**Figure 2A**). Featured UMAP plots for cell markers demonstrated that SMC, fibroblasts (FB), and macrophages (Mac) were identified as the predominant cell types in the ascending aorta (**Figure 2A-C**). While macrophages were derived solely from the nSHF lineage in both baseline and AngII-infused conditions, SMCs and fibroblasts were derived from both SHF and nSHF lineages (**Figure 2A, B**). Endothelial cells (EC) and neural cells (Neu) were also present, but the population of these cells was minimal compared to that of SMCs and fibroblasts.

**Figure 2.**
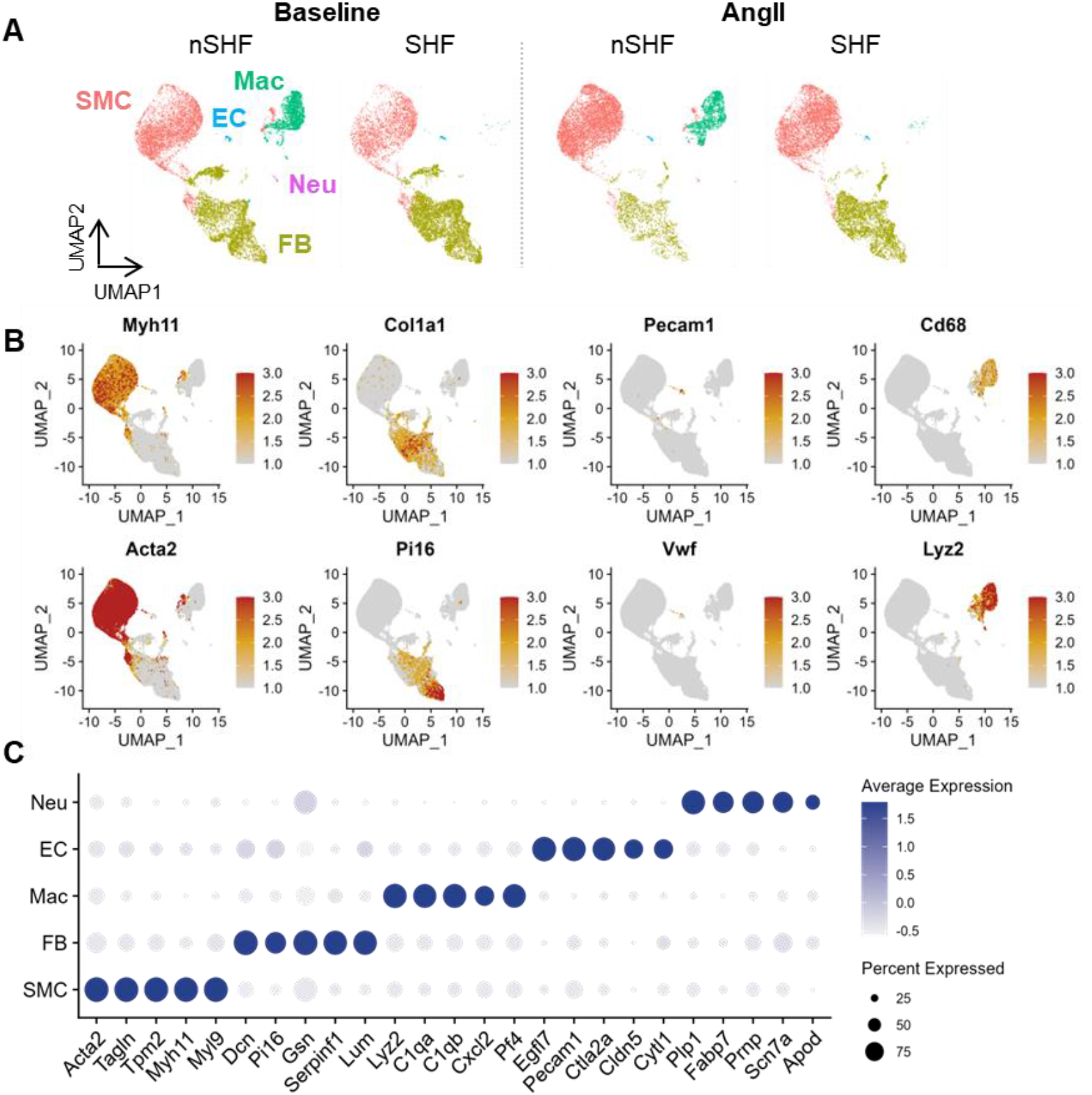
scRNAseq identified multiple cell type clusters in mouse aortas at baseline and 3 days after AngII infusion. **(A)** Uniform manifold approximation and projection (UMAP) plots of all aortic cells from nSHF and SHF origins at baseline and after 3 days of AngII infusion. SMC indicates smooth muscle cell; FB, fibroblast; EC, endothelial cell; Mac, macrophage; Neu, neural cell. **(B)** Featured UMAP plots for cell marker genes. **(C)** Dot plots for highly abundant genes in each cell type cluster.

SMC data were extracted to profile transcriptomic differences between SHF- and nSHF-origins in SMCs in response to AngII. mRNA abundance was compared between baseline and AngII in each origin separately. There were 3,381 differentially expressed genes (DEGs) identified commonly in the two origins (**Figure 3A**). Of note, 3,287 DEGs (97.2%) exhibited transcriptomic changes in the same direction in both origins (up-regulation in both origins: 47.1%, downregulation in both origins: 50.1%) and most DEGs exhibited comparable responses to AngII in SHF- and nSHF-derived SMCs.

**Figure 3.**
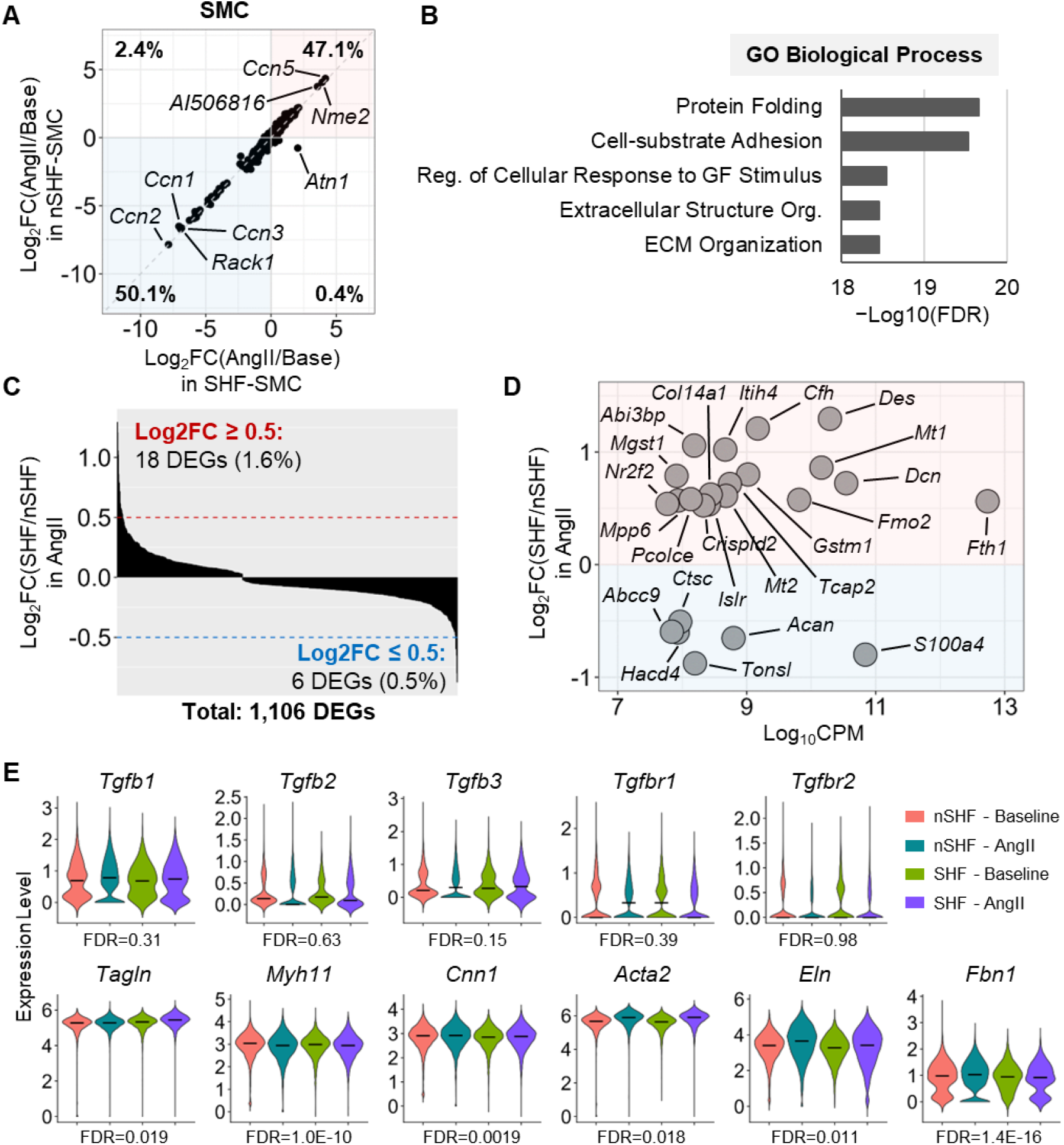
Modest transcriptomic differences between SHF- and nSHF-derived SMCs in the prepathological phase of AngII-induced thoracic aortopathy. **(A)** Dotted plot for Log_2_FC (AngII/Baseline) of DEGs in SHF- and nSHF-derived SMCs. **(B)** Top 5 annotations in GO enrichment analysis for the DEGs. **(C)** Bar plot for Log_2_FC of DEGs in SMCs. **(D)** Dotted plot for DEGs with Log_2_FC < −0.5 or > 0.5 plotted against Log_10_CPM. **(E)** Violin plots of featured DEGs in SMCs related to TGFβ, SMC contraction, and extracellular matrix. Black bars indicate the median.

Interaction analysis for AngII infusion and SMC origins identified 1,718 DEGs. Gene ontology (GO) enrichment analyses for biological processes indicated that these DEGs were primarily associated with protein folding, cell adhesion, and extracellular matrix (**Figure 3B**). Among these DEGs, 1,106 genes showed significant differences between origins during AngII infusion (**Figure 3C**). However, the fold change magnitude between origins was modest for most genes, with only 18 (1.6%) and 6 (0.5%) genes displaying Log_2_ fold changes of ≥ 0.5 or ≤ −0.5, respectively. These DEGs included *Dcn* and *Des*, which have been implicated in lineage-specific roles in the pathophysiology of aortopathy;^26,27^ however most of the DEGs lack direct evidence linking them to aortopathy (**Figure 3D**). *Mmp2* and *Mmp9*, which mediate extracellular matrix degradation, were not different between the two origins. TGF-β ligands and receptors were not statistically different in the interaction analyses (**Figure 3E**). SMC contractile genes, *Tagln, Myh11, Cnn1*, and *Acta2*, were statistically different in the interaction analyses; however, the magnitude of these gene alterations was modest. Similarly, transcriptomic differences in *Eln* and *Fbn1*, major components of elastic fibers in the aortic wall, were also modest regardless of SMC origins.

### Minor differences of transcriptomic alterations between SHF- and nSHF lineages in fibroblasts in response to short-term AngII infusion

Since fibroblasts also play a crucial role in preserving aortic integrity, we next assessed lineage-specific differences of AngII-induced transcriptomic alterations in fibroblasts. Short-term AngII infusion affected 2,900 genes in both lineages (**Figure 4A**). Similar to SMCs, most DEGs (98.8%) exhibited changes in the same direction in SHF and nSHF lineages (upregulation: 58%, downregulation: 40.8%). Two-way ANOVA interaction analysis identified 606 DEGs primarily associated with ECM (**Figure 4B, C**). However, only 11 (1.8%) and 10 (1.7%) genes exhibited Log_2_ fold changes of ≥ 0.5 or ≤ −0.5, respectively, highlighting minimal differences between the lineages (**Figure 4C**). Several molecules, such as *Il33*,^28^ *Col3a1*,^29^ and *S100A4*,^30,31^ have been reported to contribute to aortopathy formation (**Figure 4D**). Similar to SMCs, lineage-specific differences in TGF-β ligands and receptors and key elastic fiber components were either insignificant or modest (**Figure 4E**).

**Figure 4.**
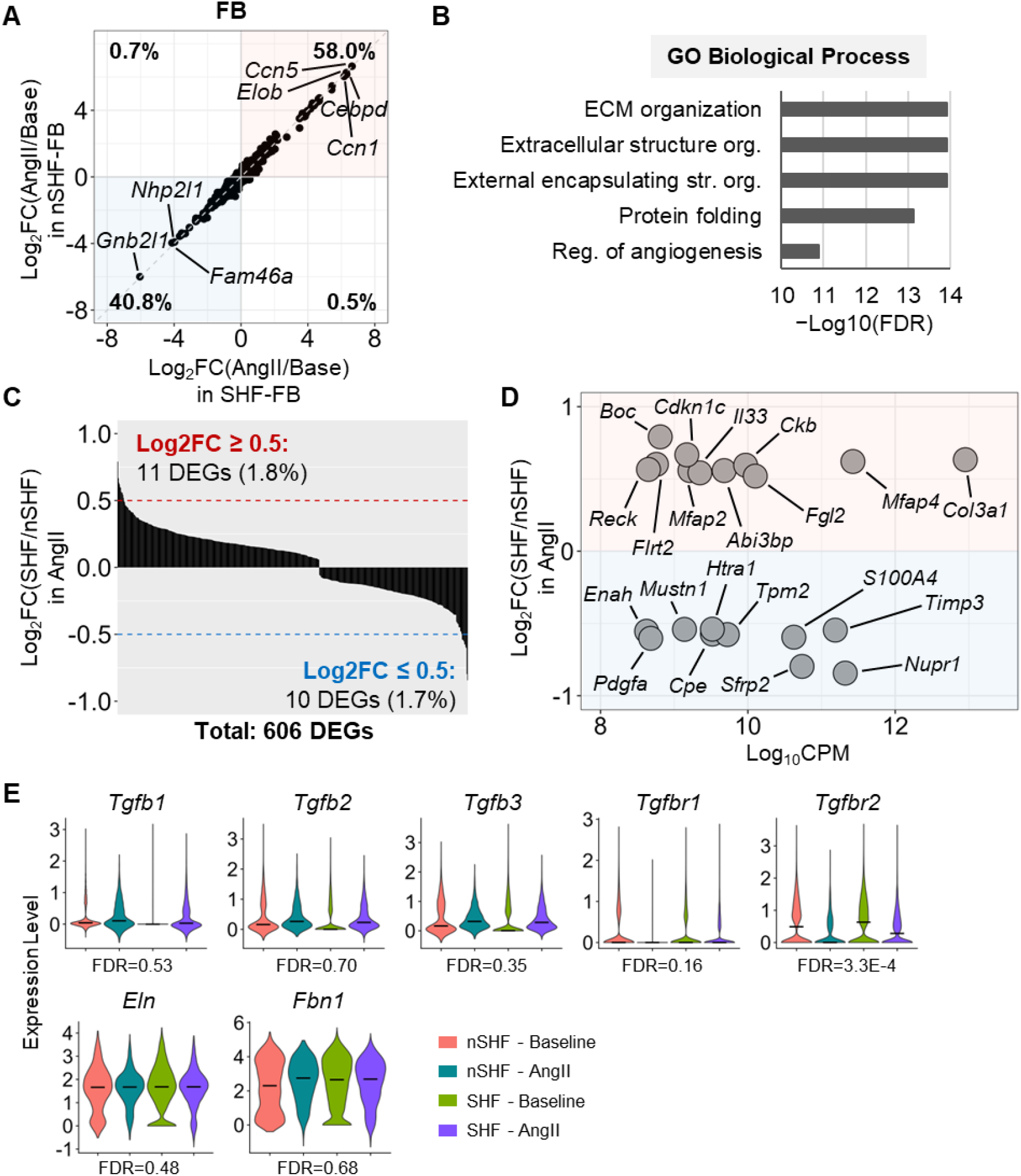
Minor differences of transcriptomic alterations between SHF- and nSHF lineages in fibroblasts in response to short-term AngII infusion. **(A)** Dotted plot for Log_2_FC (AngII/Baseline) of DEGs in SHF- and nSHF-derived FBs. **(B)** Top 5 annotations in GO enrichment analysis for the DEGs. **(C)** Bar plot for Log_2_FC of DEGs in FBs. **(D)** Dotted plot for DEGs with Log_2_FC < −0.5 or > 0.5 plotted against Log_10_CPM. **(E)** Violin plots of featured DEGs in FBs related to TGFβ and extracellular matrix. Black bars indicate the median.

### Different ECM-related transcriptomic alterations in the subcluster of FBs between SHF and nSHF origins in the prepathological phase of AngII-induced thoracic aortopathy

Our previous study identified a distinct sub-cluster in fibroblasts that may exert a deleterious role in the pathophysiology of AngII-mediated thoracic aortopathy. Therefore, we assessed potential differences regarding this fibroblast sub-cluster between SHF and nSHF lineages. In both lineages, AngII infusion developed the distinct fibroblast subcluster (**Figure 5A**, dotted red circles). This subcluster exhibited high *Pi16* abundance but lacked *Dcn* and partial expression of *Col1a* and *Ly6a* (**Figure 5B**). Our previous study identified *Ube2c* and *H2afz* as specific markers for this sub-cluster, and both markers were comparably detected in the sub-cluster of both lineages (**Figure 5C**). As fibroblasts from the distinct sub-cluster were sparse at baseline, DEG analysis was conducted between origins under AngII infusion. A total of 351 DEGs with Log_2_ fold changes of ≥ 0.5 or ≤ −0.5 were identified, including 119 upregulated and 232 downregulated genes (**Figure 5D**). Most DEGs were primarily linked to ECM processes (**Figure 5E**). Specifically, *Col15a1* and *Col18a1* were more abundant in SHF-derived cells, whereas elastin (*Eln*) and fibronectin 1 (*Fn1*), key components for maintaining elastic fiber integrity, as well as *Col3a1* and *Col8a1*, crucial for collagen fiber formation, were less abundant in the SHF-lineage fibroblast sub-cluster compared to the nSHF-lineage (**Figure 5F**). Fibroblasts in the newly emerged subcluster displayed lineage-specific differences in several ECM-related molecules in AngII-driven transcriptomic alterations, although scRNAseq analyses using all fibroblasts did not reveal obvious differences between origins.

**Figure 5.**
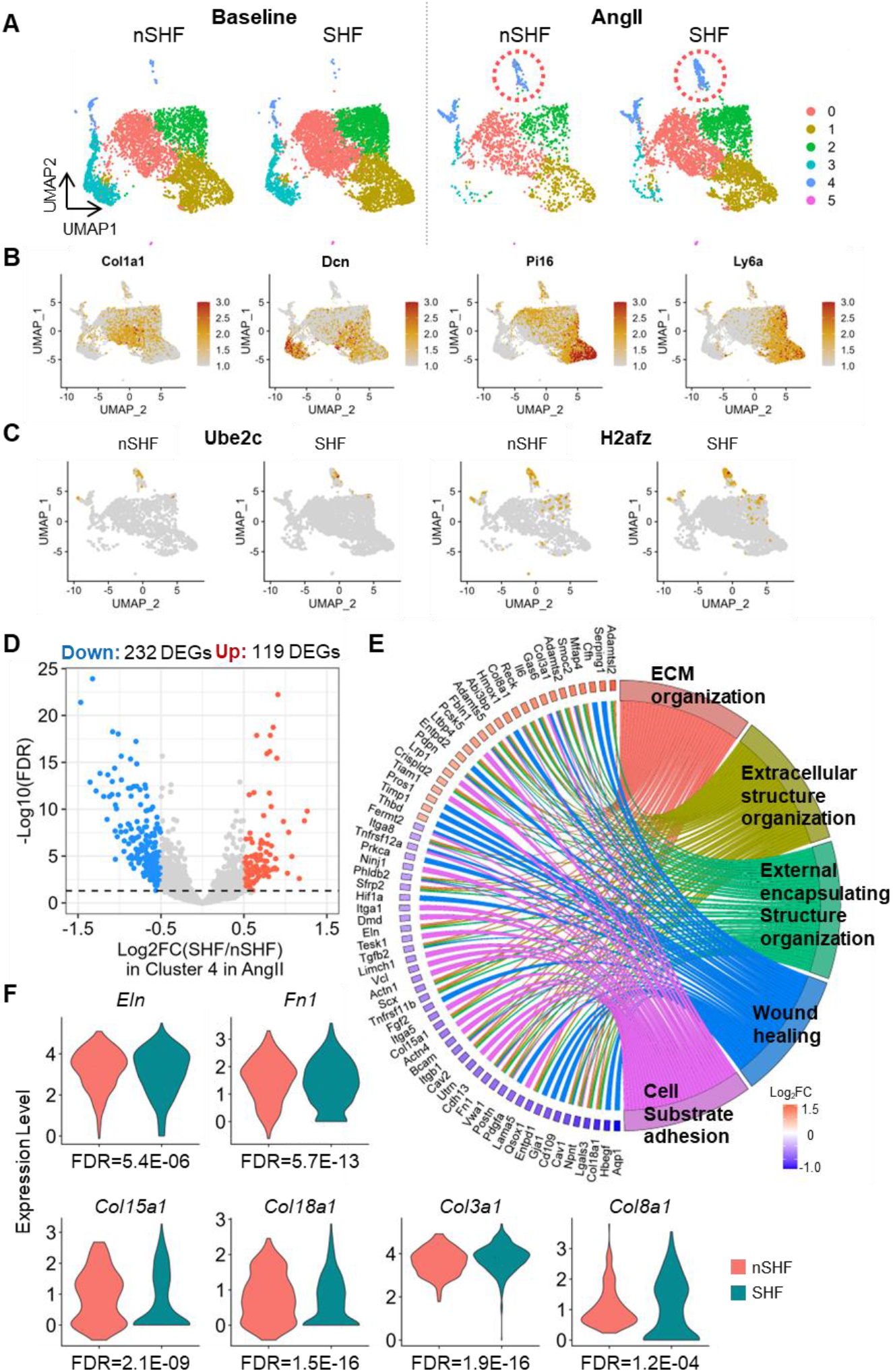
Different ECM-related transcriptomic alterations in the subcluster of FBs between SHF and nSHF origins in the prepathological phase of AngII-induced thoracic aortopathy. **(A)** Uniform manifold approximation plots (UMAP) for FBs. **(B)** Featured UMAP plots for FB marker genes, **(C)** Ube2c and H2afz. **(D)** Volcano plots and **(E)** top 5 annotations in GO enrichment analysis for DEGs of Cluster 4 in FBs. **(E)** Violin plots of featured DEGs in Culster 4 of FBs related to TGFβ and extracellular matrix.

## DISCUSSION

Our previous study demonstrated that SHF-derived SMCs are the major driver of AngII-induced thoracic aortopathy.^16^ Another group reported that AngII type 1a receptors in SHF-derived cells contributed to aortic dilatations in Loeys-Dietz syndrome model mice.^32^ These findings provide compelling evidence that SHF-derived SMCs exert a critical role in AngII-mediated thoracic aortic pathology. However, the transcriptomic profiles observed between the two origins in the present study were insignificant. In our previous study,^16^ the distributions of CNC- and SHF-derived SMCs were compared histologically in mice with AngII infusion for 4 weeks, corresponding to the chronic phase of aortopathy. The impact of SHF-specific deletion of AngII type 1a receptors was assessed at 24 weeks of age in Loeys-Dietz syndrome model mice. Thus, these studies explored the role of SHF-derived cells after development of thoracic aortopathy. In contrast, the current study investigated transcriptomic differences between origins after just 3 days of AngII infusion, prior to overt thoracic aortopathy formation, and found little transcriptomic difference between origins. These findings indicate that SHF-derived SMCs exert a critical role in progressing thoracic aortopathy, but not disease initiation. Therefore, to explore the impact of SMC origins on disease progression, it is important to examine transcriptomic differences between origins at several intervals, including the acute and advanced phases of thoracic aortopathy.

There are two previous studies that have compared transcriptomes of CNC- and SHF-derived SMCs in thoracic aortopathy formation using scRNA-seq.^26,27^ In mice with SMC-specific TGFBR1/2 deficiency, CNC- and SHF-derived SMCs displayed similar transcriptomic profiles.^27^ In contrast, in *Fbn1*^C1041G/+^ mice, there was a distinct SMC sub-cluster.^26^ Interestingly, several ECM-related genes, including *Col1a1* and *Col3a1*, were upregulated in SHF-derived SMCs compared to non-SHF-derived SMCs of the distinct sub-cluster.^26^ Of note, our previous and present studies identified a distinct sub-cluster only in fibroblasts, but not in SMCs. In addition, we found a decrease in *Col3a1* mRNA abundance in SHF-derived fibroblasts of a unique cluster. Thus, inconsistencies remain regarding the existence of SMC sub-populations and ECM-related gene profiles across studies. Potential explanations for these differences could be the use of different mouse models for thoracic aortopathy and different promoters for SHF-lineage labeling. Our study used the AngII-induced thoracic aortopathy model, while previous studies used *Fbn1*^C1041G/+^ and SMC-specific TGFBR1/2 deficient mice, both of which are driven by the dysregulation of TGF-β signaling. In addition, while previous studies used the Nkx2.5 promoter,^26,27^ we used the Mef2c promoter. Although both Nkx2.5 and Mef2c promoters are commonly used for tracking SHF-derived cells in cardiovascular tissue, these promoters have disparities in their distributions.^33^ Therefore, it is possible that the difference in models and promoters is attributed to induce inconsistencies across the studies.

In the present study, scRNAseq identified a distinct sub-cluster in fibroblasts originating from both SHF- and nSHF lineages prior to the occurrence of thoracic aortopathy. Of note, fibroblasts in the newly identified subcluster exhibited lineage-specific alterations in mRNA abundance of important ECM genes, including *Eln* and *Col3a1*, following short-term AngII infusion. Genetic mutations in *Eln* and *Col3a1* have been identified in patients with Williams syndrome^34^ and vascular-type Ehlers-Danlos syndrome^35^, respectively, both of which are associated with aortic abnormalities in humans. Mouse models with genetic deletions and mutations in *Eln* and *Col3a1* recapitulate aortic manifestations of these syndromes.^29,36^ Therefore, the downregulation of *Eln* and *Col3a1* in selected fibroblasts from the SHF origin may play a critical role in compromising aortic integrity by AngII infusion. However, these molecules were not significantly different when analyzing the entire fibroblast population. Given the low number of newly emerged fibroblasts at 3 days of AngII infusion, this is likely due to the dilution of the differences by other fibroblasts. It is possible that this distinct fibroblast population may expand with longer exposure and disease progression. Since our previous and current studies identified *Ube2c* and *H2afz* as specific markers for this unique fibroblast subcluster, further study using these markers could provide deeper insights into fibroblast heterogeneity and its role in the pathophysiology of thoracic aortopathy.

The present study used Mef2c-Cre +/0 R26R^mTmG/0^ mice to trace SHF-derived cells; mGFP protein is expressed in cells driven by a Mef2c-Cre promoter, which enabled the selection of SHF-derived cells. Conversely, mTomato-positive cells originate from other sources, including the CNC. Since SMCs in the ascending aorta are from both SHF and CNC origin, most non-SHF-derived cells are considered to be derived from the CNC. However, several studies have reported that adventitial plastic cells, whose embryonic origin has not been determined, contribute to pathological remodeling of the media.^37^ These cells may interfere with the interpretation of a comparison between SHF-vs CNC-derived cells. Further study is needed to directly determine transcriptomic alteration in CNC-derived SMCs using Wnt1-Cre, a CNC-specific promotor.

In conclusion, although selected fibroblasts exhibited lineage-specific differences in several ECM-related molecules, entire SMC and fibroblast transcriptomes exhibit modest differences between SHF-derived cells and those from other origins during the prepathological phase of AngII-induced thoracic aortopathy.

## Supplemental Materials

Supplemental Figure 1 - 3

Supplemental Excel File

---

**Supplementary Figure 1.**
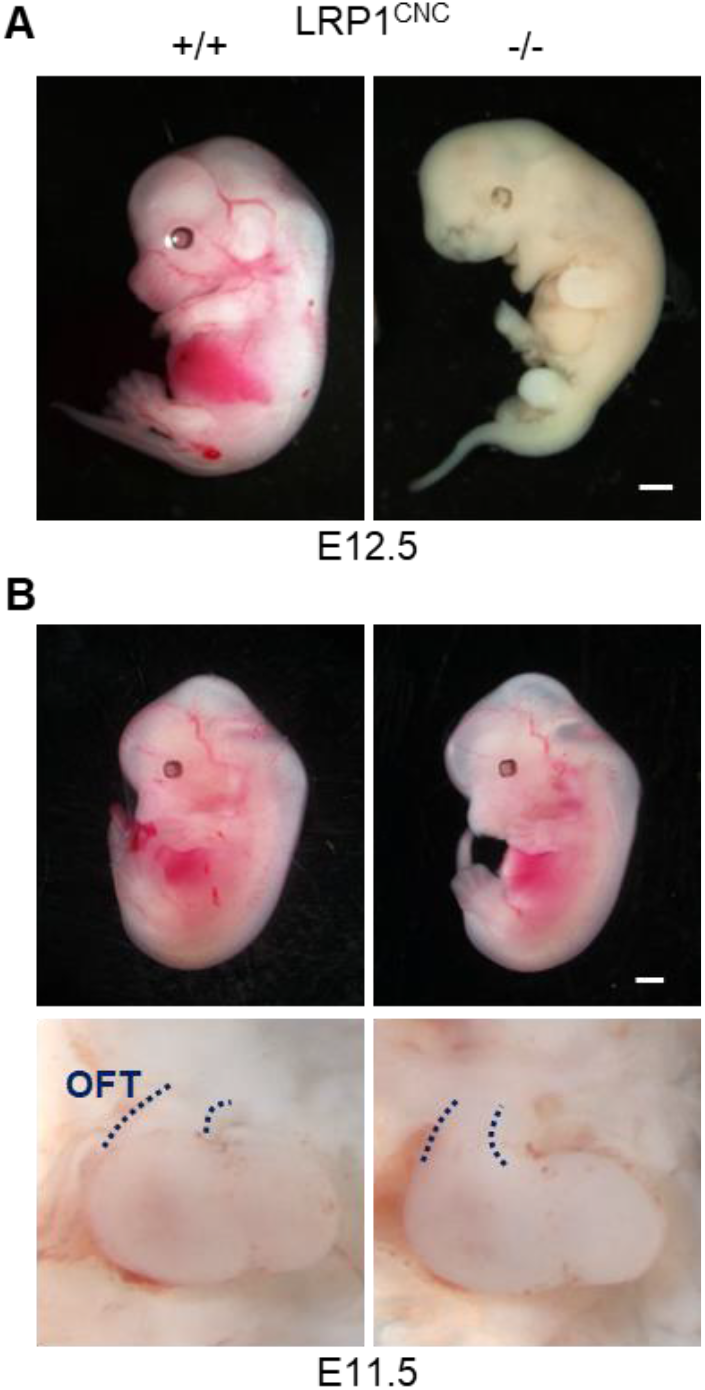
Embryonic lethality provoked by LRP1 deletion in CNC-derived cells. **(A)** Gross appearance of wild type and CNC-specific LRP1-deleted fetuses at E12.5 (n = 5 to 7 per group). **(B)** Gross appearance of fetuses and outflow tracts in wild type and CNC-specific LRP1-deleted fetuses at E11.5 (n = 5 per group). Blue dotted lines indicate the outflow tract (OFT). Scale bar = 1 mm.

**Supplementary Figure 2.**
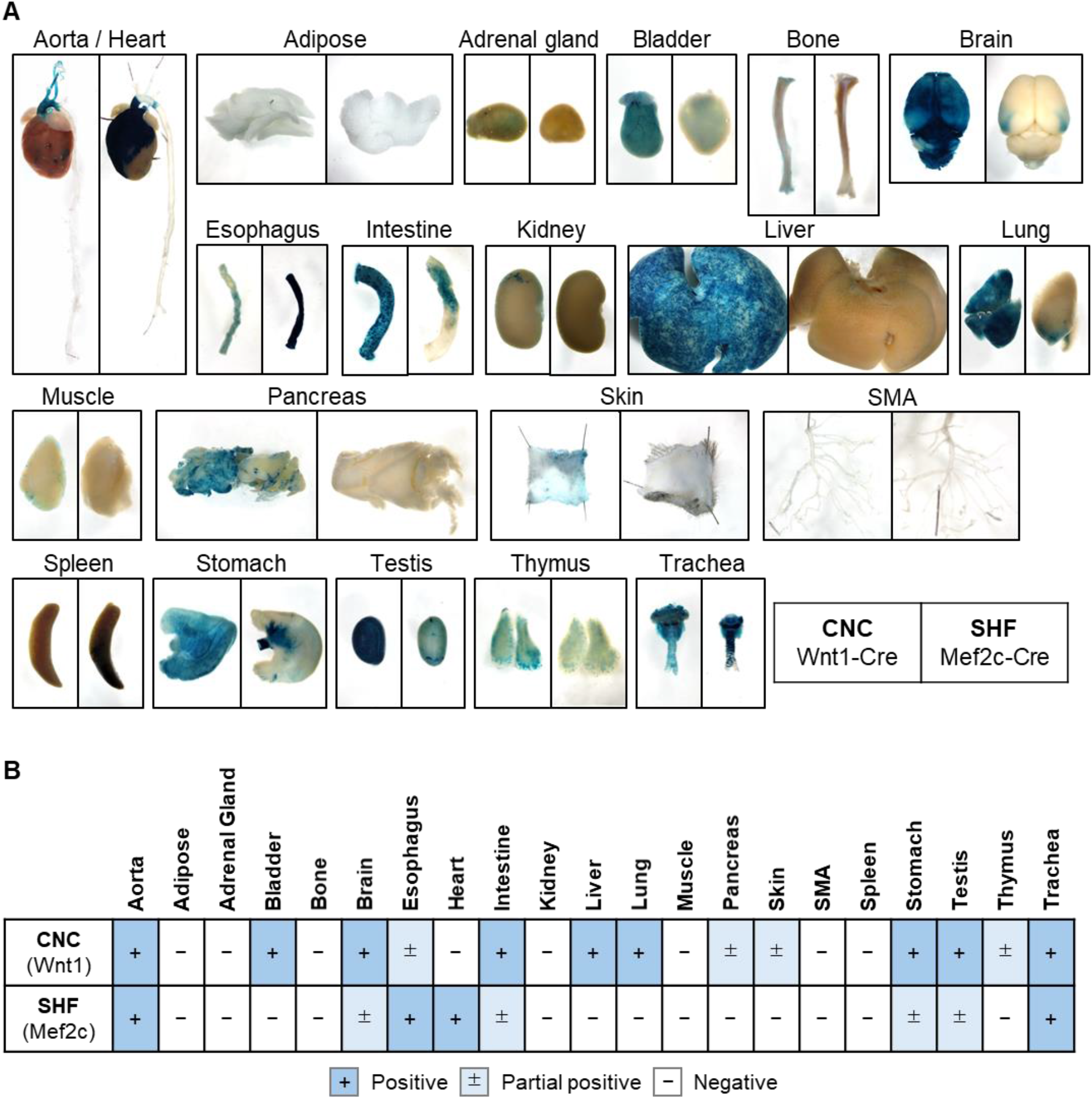
Distinct distribution of CNC- and SHF-derived cells in major organs in mice. Representative **(A)** gross images of X-gal staining of major organs in male Wnt1- and Mef2c-Cre +/0 R26R^Lacz^ +/0 mice and **(B)** its summary table (n = 3 per group). SMA indicates superior mesenteric artery.

**Supplementary Figure 3.**
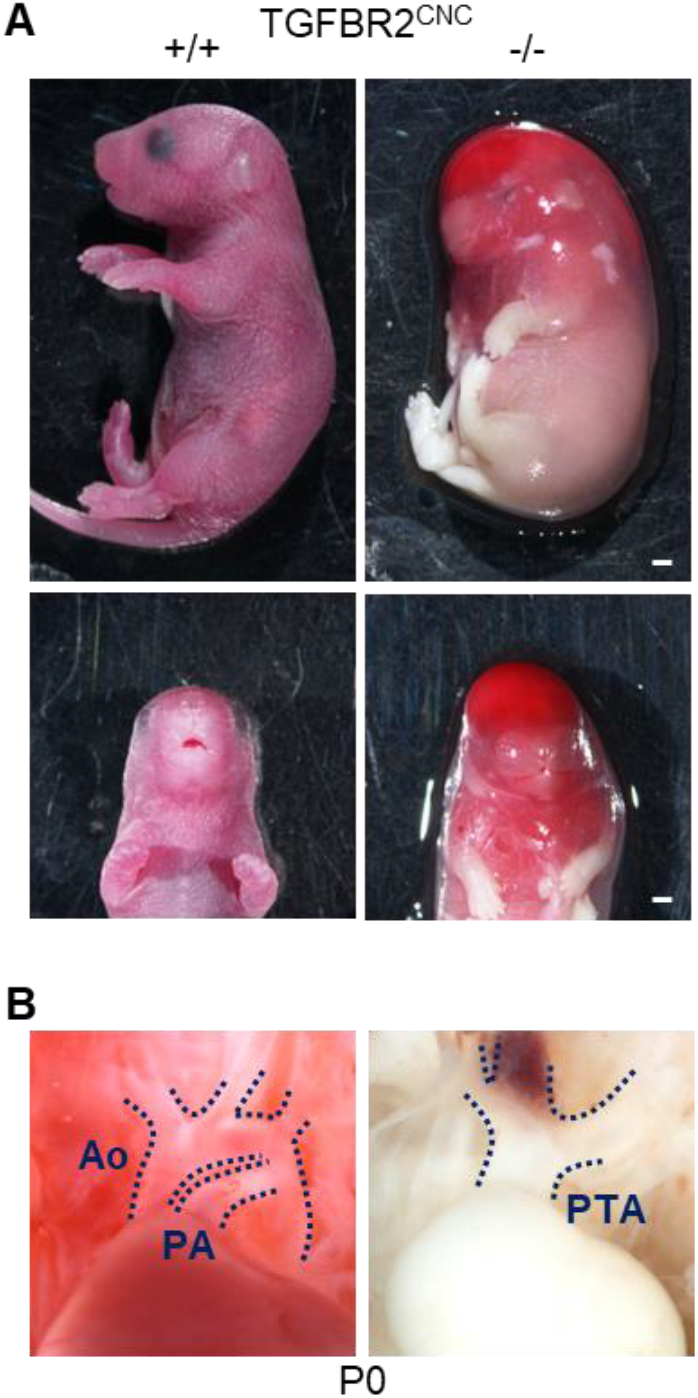
Calvaria defect and persistent truncus arteriosus in mice with TGFBR2 deletion in CNC-derived cells. **(A)** Gross appearance and **(B)** in situ images of the thoracic aorta in wild-type and CNC-specific TGFBR2-deleted neonates at P0 (n = 3 to 5 per group). Blue dotted lines indicate the aorta, (Ao), pulmonary artery (PA), and persistent truncus arteriosus (PTA). Scale bar = 1 mm.

